# *Staphylococcus aureus* Phenol-Soluble Modulins Mediate Interspecies Competition with Upper Respiratory Commensal Bacteria

**DOI:** 10.1101/2024.09.24.614779

**Authors:** Joshua T Huffines, Megan R Kiedrowski

**Author notes:** Address correspondence to Megan Kiedrowski, **Email:**.

## Abstract

In chronic rhinosinusitis (CRS) disease, microbial dysbiosis is considered a key contributor to inflammation and pathogenicity, with increased prevalence of upper respiratory tract (URT) pathogens concomitant with decreased abundance of commensal species. *Staphylococcus aureus* is a common URT pathobiont associated with higher carriage rates in CRS. *S. aureus* secreted toxins are implicated in CRS pathogenesis, and toxins and antibodies to *S. aureus* secreted factors have been observed in tissue from CRS subjects. CRS disease severity is positively correlated with immune reactivity to *S. aureus* proteins. Prior studies have examined polymicrobial interactions between *S. aureus* and URT commensals, however, no studies to date have described possible methods employed by *S. aureus* to outcompete commensals leading to a *S. aureus-*dominant microbiome as seen in CRS. This study addresses this gap in knowledge by characterizing how a CRS-associated secreted toxin from *S. aureus* can inhibit aggregation in commensal URT species. Using a model URT commensal, *Corynebacterium pseudodiphtheriticum*, we identified a CRS-associated secreted protein from *S. aureus*, δ-toxin (Hld), that can inhibit *C. pseudodiphtheriticum* aggregation at biologically relevant concentrations. Furthermore, we observed recombinant δ-toxin reduces *C. pseudodiphtheriticum* adherence and aggregation on human nasal epithelial cells in an air-liquid interface cell culture model. These results define a novel mechanism by which *S. aureus* can disrupt URT commensal lifestyles of microbial competitors, contributing to the establishment of microbial dysbiosis.

**IMPORTANCE:** Microbial dysbiosis in the upper respiratory tract (URT) is associated with disease pathogenicity in chronic rhinosinusitis (CRS). There are significant links between *Staphylococcus aureus* and worse CRS outcomes, but no studies to date have demonstrated if *S. aureus* outcompetes other URT microbes through direct interactions. Here, we report that *S. aureus* δ-toxin, a secreted protein found in CRS patient tissue, can inhibit the ability of commensal bacteria to aggregate, adhere to, and grow in association with human nasal epithelial cells. These results suggest a potential mechanism for *S. aureus* to establish dominance in the URT microbiome through direct antagonism of commensals with a disease-associated toxin.

## INTRODUCTION

Chronic rhinosinusitis (CRS) is an inflammatory disease of the upper respiratory tract (URT) with high prevalence in the United States leading to annual costs approaching $19 billion (1–3). Microbial dysbiosis is considered a significant contributor to CRS pathogenicity, with a shift from highly abundant keystone commensal species to URT bacterial pathogens (4–6). Among these pathogens, *Staphylococcus aureus* is significantly associated with CRS pathogenicity. Increased *S. aureus* carriage in CRS patients is correlated with disease severity and recurrence, toxin presence in tissue samples, and antibody specificity to well-known *S. aureus* virulence factors (7–10).

Many *S. aureus* CRS-associated toxins require activation of the accessory gene regulator (agr) quorum-sensing system (7, 11) to upregulate their production. The agr system governs the switch between the staphylococcal biofilm lifestyle and acute virulence, with agr deactivation or inhibition being associated with commensalism and reduced virulence (12, 13). When activated through sensing of the *S. aureus* auto-inducing peptide (AIP) by its cognate membrane-localized histidine kinase, AgrC, phosphorylation of the response regulator AgrA leads to transcription of RNAII, encoding the agr machinery, and RNAIII, the primary downstream effector that governs toxin production (11).

The *S. aureus* agr system is involved in many of the polymicrobial interactions between *S. aureus* and other URT microbes, with common, highly abundant commensal species inhibiting or exploiting agr to limit *S. aureus* growth and virulence (14–16). Previous work on polymicrobial interactions between *S. aureus* and URT commensals has largely focused on methods by which commensal species can inhibit or limit *S. aureus* (17, 18). Considering the central role of agr in both CRS pathogenicity and polymicrobial interactions, a gap in knowledge exists for how *S. aureus* can overcome this inhibition to outcompete URT commensals and to become dominant in the CRS microbiome in sinus disease. The *Corynebacterium* genus contains many species considered to be commensals of the URT that can be found colonizing healthy individuals in high abundance. *Corynebacterium* has been found to be negatively correlated with *S. aureus* carriage in URT microbiome analyses (19, 20). Other groups have examined possible mechanisms for how *Corynebacterium* spp. can contribute to decreased *S. aureus* carriage in the healthy URT (15, 16). Previously, we examined interactions between *Corynebacterium* with *S. aureus* using an air-liquid interface cell culture model (21). Using a sequential model of infection, we observed *S. aureus* was capable of colonizing human nasal epithelial cells (HNECs) despite the presence of *C. pseudodiphtheriticum*. Further, co-colonization with *S. aureus* led to an observable decrease in *C. pseudodiphtheriticum* aggregate formation on HNECs. This suggested to us that in CRS disease, where *S. aureus* is highly abundant in the inflamed URT, *S. aureus* may be capable of outcompeting *Corynebacterium* occupying the same niche.

Here, we identify a novel function for a CRS-associated *S. aureus* secreted toxin that can directly alter *Corynebacterium* aggregation and adherence to human nasal epithelial cells. Utilizing a microscopy-based aggregation assay with a model commensal bacterium commonly found in the healthy URT, *Corynebacterium pseudodiphtheriticum*, we found*S. aureus* secretions can inhibit *C. pseudodiphtheriticum* aggregation in a non-bactericidal manner. Next, utilizing the *S. aureus* Nebraska Transposon Mutant Library (22), we identified the secreted factor from *S. aureus* to be agr-regulated and dependent on the phenol-soluble modulin (PSM) transporter (PMT) complex. Further testing of *S. aureus* mutants lacking PSM genes, as well as addition of purified PSM peptides, confirmed the ability of PSMs to inhibit *C. pseudodiphtheriticum* aggregation and identified δ-toxin as the primary PSM responsible for this activity. Using an air-liquid interface nasal epithelial cell culture model to evaluate *C. pseudodiphtheriticum* colonization (21), we demonstrate the ability of exogenous δ-toxin to inhibit *C. pseudodiphtheriticum* adherence to the nasal epithelium. Taken together, this study characterizes a mechanism by which *S. aureus* can inhibit URT commensals with a CRS-associated toxin, contributing to the microbial dysbiosis commonly observed in CRS disease.

## RESULTS

### Staphylococcus aureus secreted factors inhibit Corynebacterium aggregation

We recently observed *Corynebacterium* sinus isolates, including *C. pseudodiphtheriticum*, grew more slowly in vitro in comparison to *S. aureus* (21). Therefore, to test if *S. aureus* affects commensal bacterial growth or behavior independently of competition for nutrients in co-culture, we first examined the effects of *S. aureus* secreted factors on *C. pseudodiphtheriticum* growth as a model URT commensal species using cell-free conditioned medium (*Sa* CFCM) prepared from stationary-phase *S. aureus* cultures. Using a tdTomato-expressing *C. pseudodiphtheriticum* strain (21), we observed *C. pseudodiphtheriticum* forms large aggregates in liquid culture, however there was a substantial decrease in the size of *C. pseudodiphtheriticum* aggregates when grown in the presence of *Sa* CFCM (Fig. 1A). We developed a high-throughput, quantifiable microscopy-based aggregation assay to further characterize this phenotype (Supplementary Methods). Average *C. pseudodiphtheriticum* aggregate size was significantly reduced when grown in the presence of *Sa* CFCM, even at levels below 5% v/v (Fig. 1B). *Sa* CFCM did not significantly impact *C. pseudodiphtheriticum* growth, even when added at up to 25% v/v to *C. pseudodiphtheriticum* cultures (Fig. 1C). *C. pseudodiphtheriticum* colony forming units were only observed to decrease substantially from untreated control cultures in the presence of 50% v/v *Sa* CFCM, likely attributable to lack of nutrients. Using live imaging, we assessed whether *C. pseudodiphtheriticum* was actively growing as aggregates and found that log_2_ transformed mean aggregate area steadily increased at a linear rate over 20 hours (Fig. S1; Movie S1).

**Figure 1:**
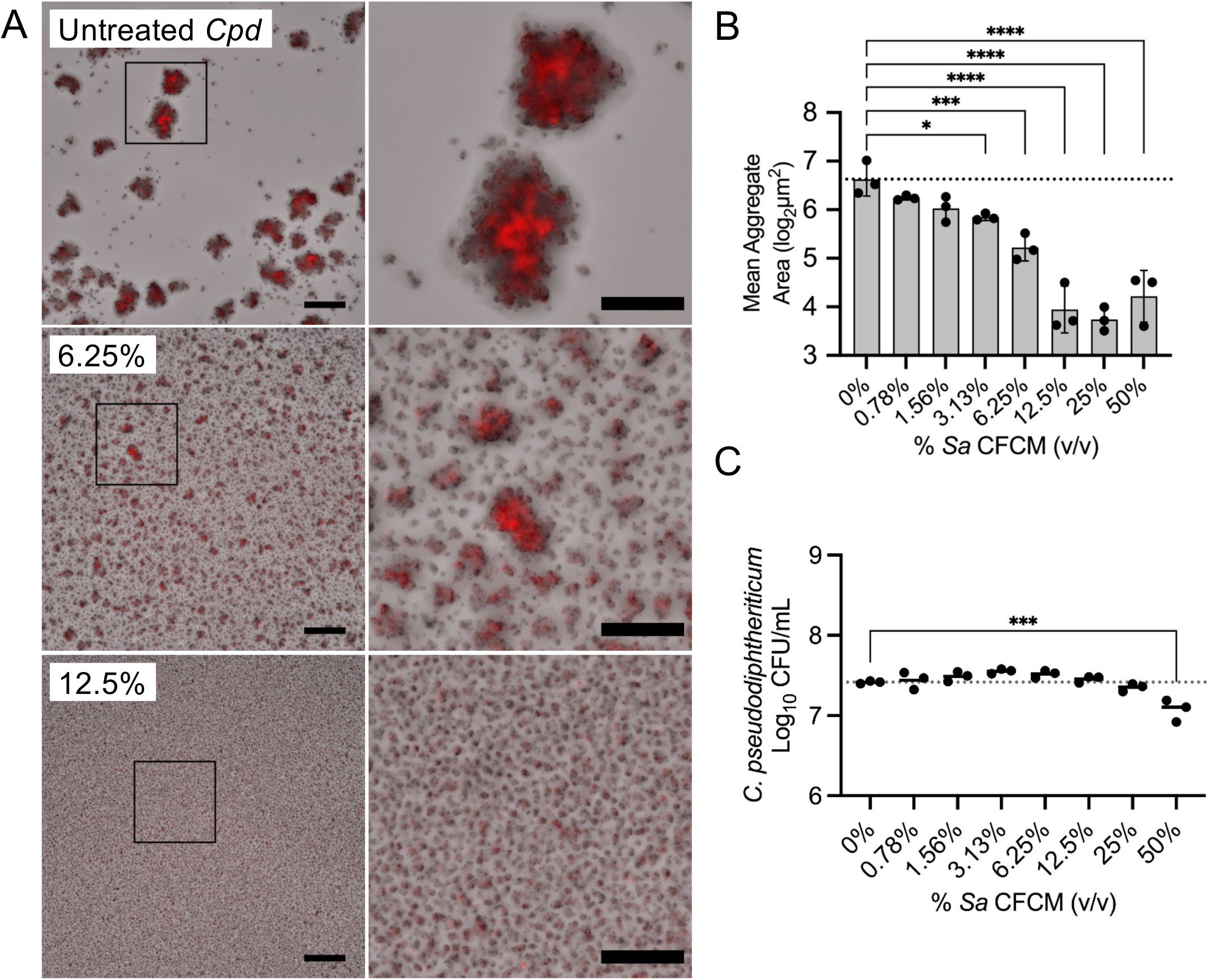
Staphylococcus aureus secreted factors inhibit Corynebacterium pseudodiphtheriticum aggregate formation. (A) tdTomato *C. pseudodiphtheriticum* grown in brain-heart infusion broth (BHI) for 20 hours with increasing concentrations of *S. aureus* CFCM prepared from *S. aureus* cultures grown for 24 hours at 37°C. 8X magnification of inset to right. Images are representative of 3 biological replicates with 3-6 technical replicates each. Scale = 80µm, 40µm for 8X magnification. (B) Mean aggregate size for images in A. *n*=3 biological replicates with 3-6 technical replicates each. (C) Colony-forming units paired with A and B. *n*=3 biological replicates with 3 technical replicates each. Significance determined using a one-way ANOVA with Dunnett’s multiple comparisons test. \*\**P*<0.01, \*\*\**P*<0.001, \*\*\*\**P*<0.0001.

To determine whether production of secreted factors capable of inhibiting *C. pseudodiphtheriticum* aggregate formation was broadly conserved in *S. aureus*, we screened CFCM from several common *S. aureus* lab strains. We observed variation in anti-aggregation ability amongst the seven isolates evaluated, with strains SH1000, 502A and USA300 SF8300 having similar levels of activity as USA300 LAC (Fig. 2). In the presence of CFCM from some *S. aureus* strains, including a USA100 MRSA isolate, Newman, N315 and Mu50, *C. pseudodiphtheriticum* aggregation did not differ significantly from the untreated control. In line with results using CFCM from *S. aureus* strain USA300 LAC 13c (Fig. 1C), we did not observe large differences in *C. pseudodiphtheriticum* viability in the presence of CFCM at 25% v/v from additional *S. aureus* strains tested, with the exception of *S. aureus* Newman (Fig. 2C).

**Figure 2:**
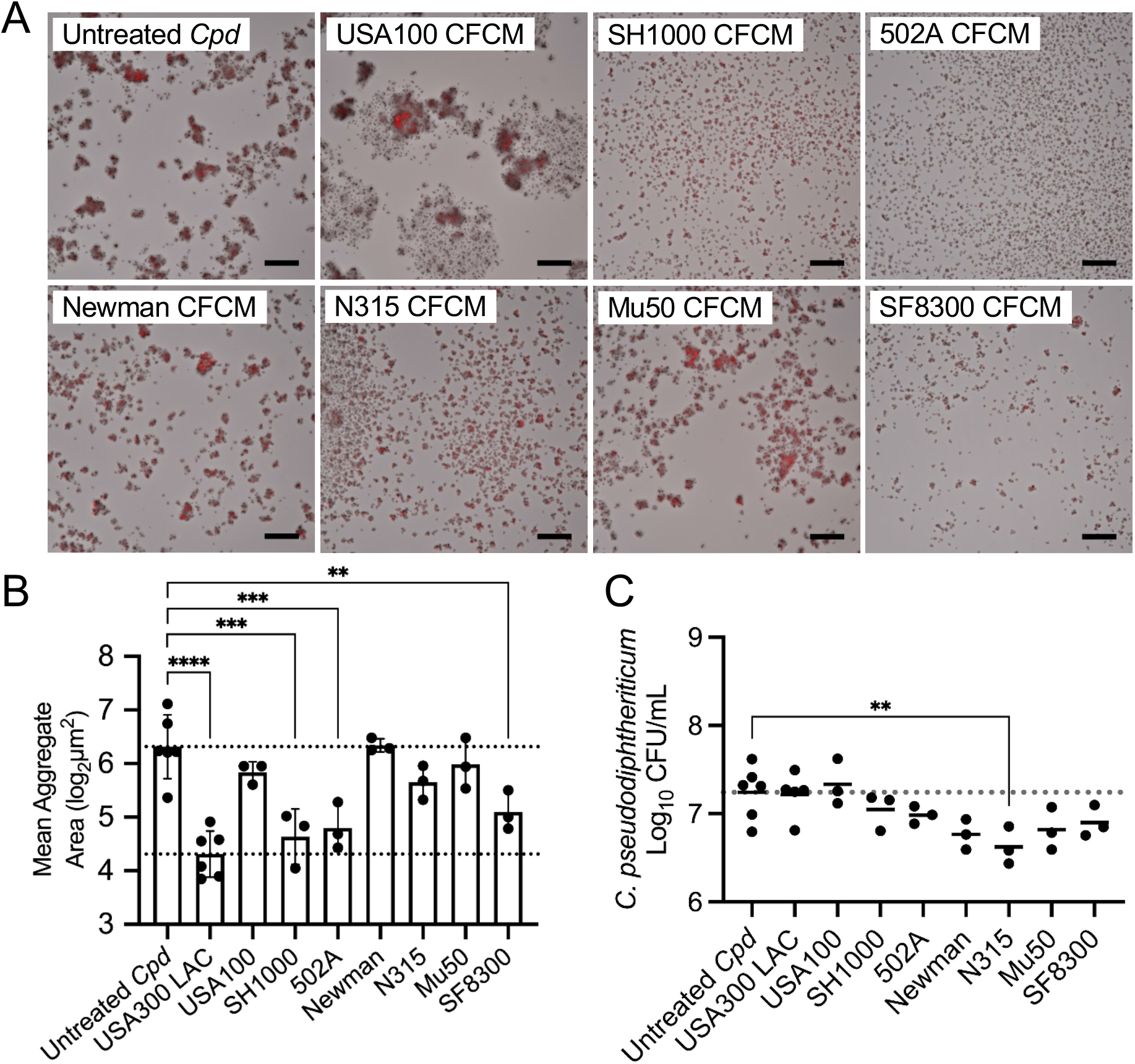
Anti-aggregation activity of *Staphylococcus aureus* secreted factors towards *Corynebacterium pseudodiphtheriticum* varies across strains. (A) tdTomato *C. pseudodiphtheriticum* grown for 20 hours in BHI broth with 25% BHI or *S. aureus* CFCM prepared from *S. aureus* cultures grown for 24 hours at 37°C. n = 3 biological replicates with 3-6 technical replicates each. (B) Quantification of mean aggregate area from images shown in A. (C) Colony-forming units of *C. pseudodiphtheriticum* from A. *n*=3-6 biological replicates with 3 technical replicates each. Significance determined by one-way ANOVA with Dunnett’s multiple comparisons test. \*\**P*<0.01, \*\*\**P*<0.001, \*\*\*\**P*<0.0001.

We next examined whether the ability of *Sa* CFCM to prevent *Corynebacterium* aggregation applied to other *Corynebacterium* species and strains (Fig. 3). Microscopy showed a CRS clinical isolate of *Corynebacterium propinquum*, an ATCC strain of *Corynebacterium pseudodiphtheriticum*, and a skin isolate of *Corynebacterium amycolatum* all formed large aggregates when cultured in BHI broth, whereas a *Corynebacterium accolens* isolate formed smaller aggregates of a substantially smaller area (Fig. 3A,B). The three strains that formed large aggregates had significantly lower average aggregate area when grown in the presence of 10% v/v *Sa* CFCM, similar to what we observed with our model CRS *C. pseudodiphtheriticum* strain (Fig. 3B). These strains also showed no defect in growth in the presence of *Sa* CFCM as measured by colony forming unit counts (Fig. 3C). However, average aggregate size of *C. accolens* was not significantly affected by *Sa* CFCM, likely attributable to the comparatively small aggregate size in the untreated group (Fig. 3A,B).

**Figure 3:**
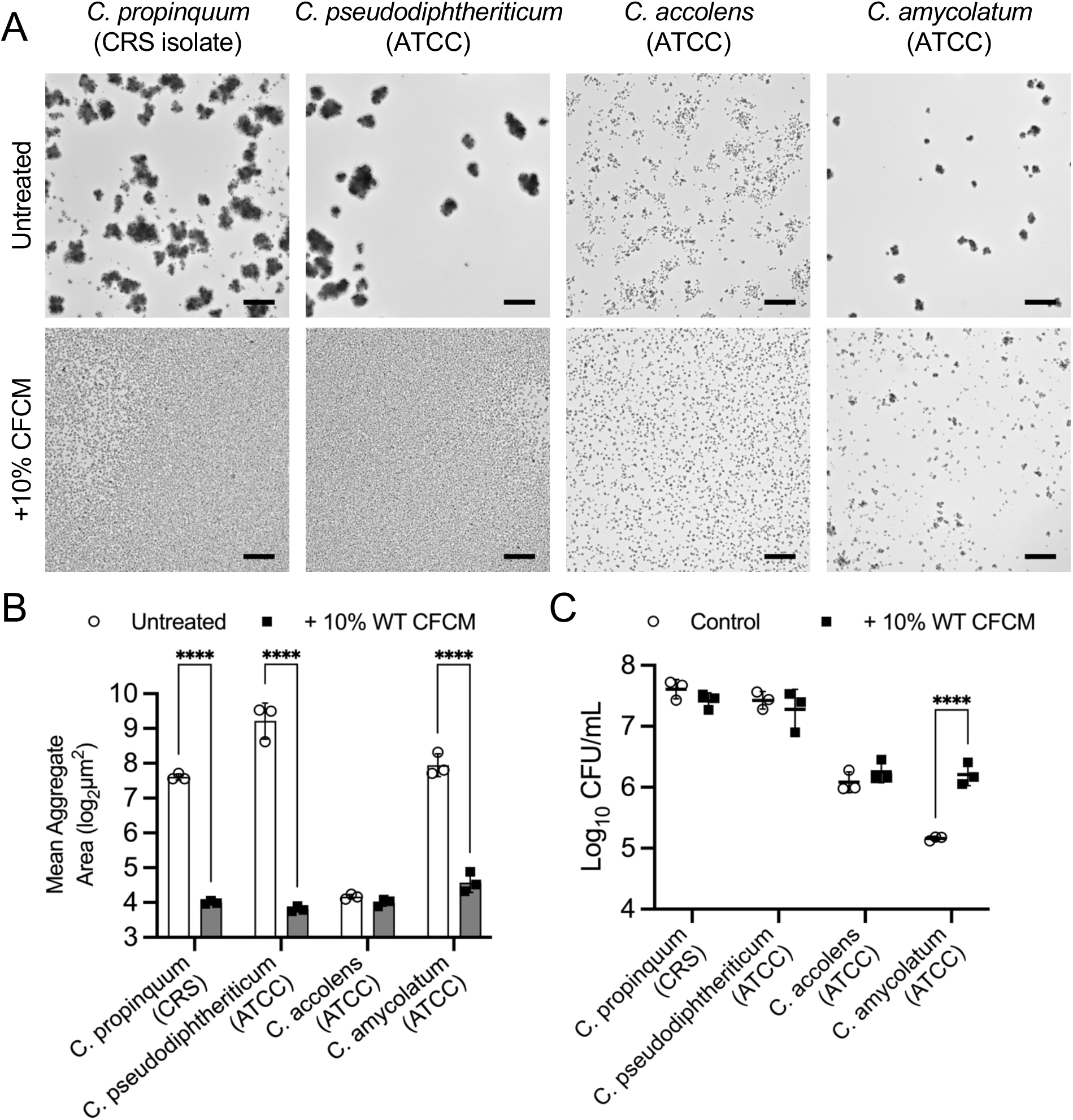
*Staphylococcus aureus* secreted factors inhibit aggregation of multiple *Corynebacterium* species. (A) *Corynebacterium* species grown for 16 hours in BHI broth with 10% BHI or *S. aureus* CFCM prepared from *S. aureus* cultures grown for 24 hours at 37°C. Images are representative of 3 biological replicates with 6 technical replicates each. (B) Quantification of mean aggregate area from images shown in A. Significance determined by two-way ANOVA with Sidak multiple comparisons test. \*\*\**P*<0.001, \*\*\*\**P*<0.0001.

### Secreted proteins from Staphylococcus aureus inhibit Corynebacterium pseudodiphtheriticum aggregation

To characterize the nature of the *S. aureus* secreted factor preventing *C. pseudodiptheriticum* aggregation, we performed a series of treatments on *S. aureus* USA300 CFCM and prior to testing treated CFCM in our aggregation assay at the highest percentage that did not significantly reduce growth (Fig. 4). Heat-treated *Sa* CFCM retained the ability to inhibit *C. pseudodiphtheriticum* aggregation and reduce aggregate size. Hypothesizing that *S. aureus*-derived metabolites or small molecules could be influencing *C. pseudodiphtheriticum* aggregation, we tested *Sa* CFCM passed through an Amicon centrifugal filter with a 3 kilodalton cutoff and found aggregate size was no longer affected, resembling untreated *C. pseudodiphtheriticum*. Addition of the *Sa* CFCM fraction retained in the centrifugal filter (diluted to the original volume to limit effects of concentration) led to significant reduction in aggregate size, as observed in the presence of untreated *Sa* CFCM. Given this, we hypothesized a secreted, heat-resistant *S. aureus* protein mediates aggregate inhibition. *Sa* CFCM treated with proteinase-K followed by heat-inactivation lost the ability to inhibit *C. pseudodiphtheriticum* aggregation, resulting in aggregates similar in size to a media-only untreated control.

**Figure 4:**
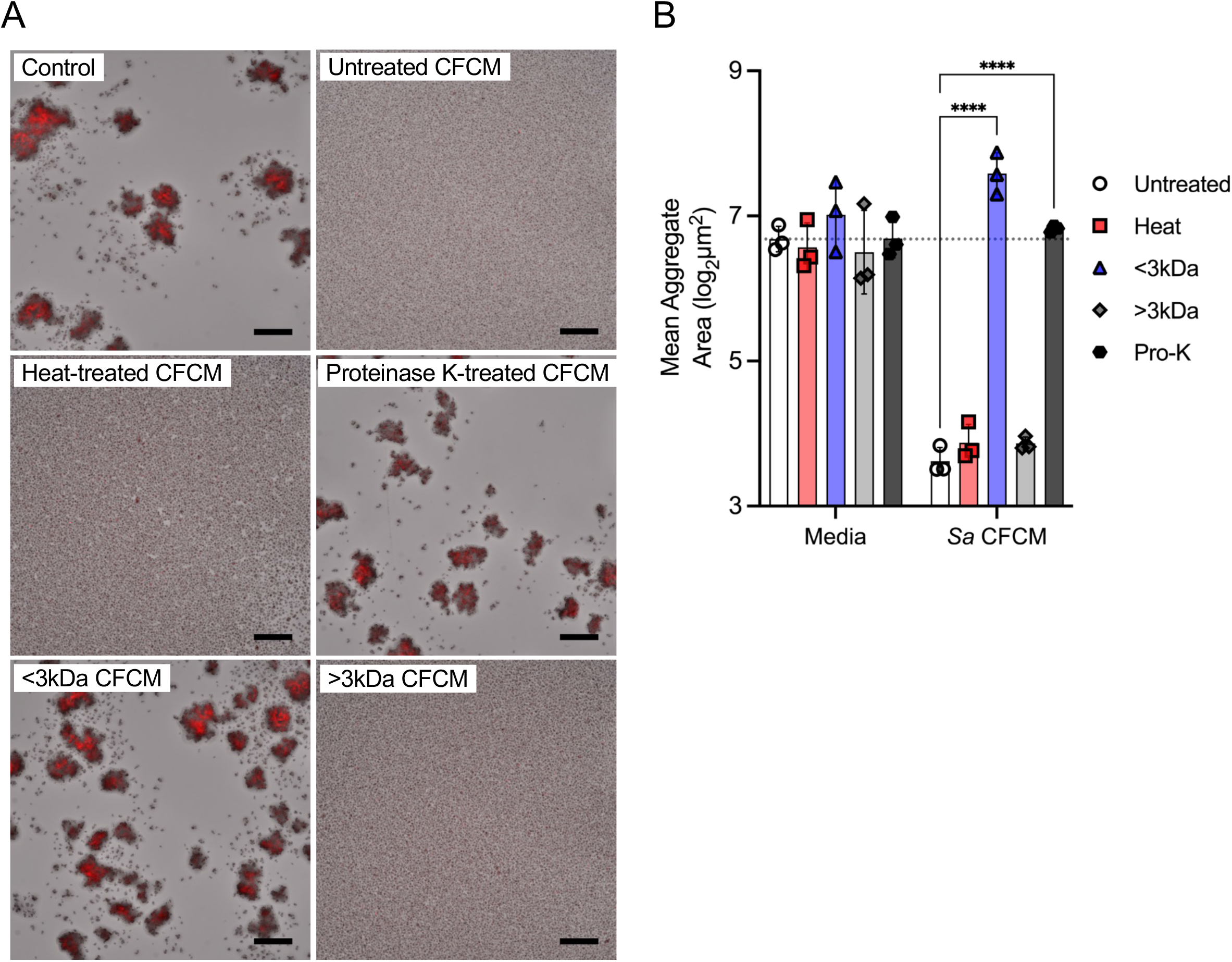
Staphylococcus aureus secreted proteins inhibit Corynebacterium pseudodiphtheriticum aggregate formation. (A) tdTomato *C. pseudodiphtheriticum* grown for 20 hours in BHI broth with 25% BHI or *S. aureus* CFCM that was untreated, heated at 98°C for 1 hour, incubated with 100 µg/mL proteinase-K, filtered using a 3kDa-cutoff Amicon filter, or concentrated in the filter and diluted to the original concentration. *n*=3 biological replicates with 3 technical replicates each. Scale bar = 80µm. (B) Mean aggregation size for images in A. *n*=3 biological replicates with 3 technical replicates each. Significance determined by two-way ANOVA with Dunnett’s multiple comparisons test (\*\*\*\**P*<0.0001).

### Agr-regulated phenol-soluble modulins from *Staphylococcus aureus* inhibit *C. pseudodiphtheriticum* aggregation

Results of CFCM treatments suggested the secreted *S. aureus* anti-aggregation factor capable of targeting other commensal species was a heat-stable protein larger than 3 kDa. We next utilized the Nebraska Transposon Mutant Library (22) to conduct a targeted screen to identify genes involved in the regulation, production, and secretion of this factor (Fig. 5). We began by screening mutants in well-characterized regulatory genes and identified expression of *agrA*, encoding the response regulator for the agr quorum-sensing system in *S. aureus*, and *sarA* encoding a positive regulator for agr quorum-sensing (23), were necessary to inhibit *C. pseudodiphtheriticum* aggregation (Fig. 5; blue bars).

**Figure 5:**
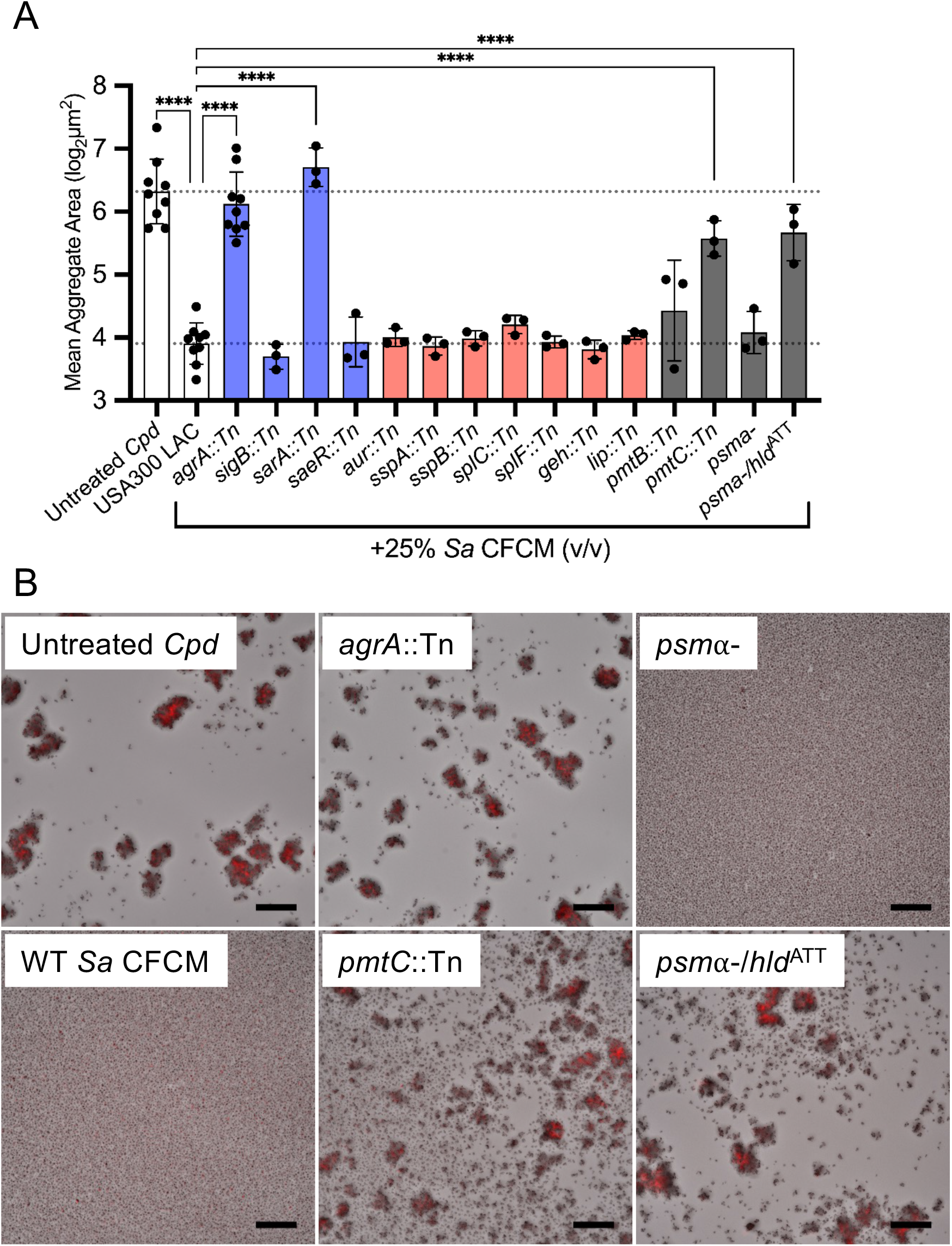
S. aureus hld is required to inhibit Corynebacterium pseudodiphtheriticum aggregation. (A) Mean aggregate size quantification of tdTomato *C. pseudodiphtheriticum* exposed to 25% *S. aureus* CFCM from USA300 LAC 13, transposon insertion mutants, or PSM-deficient strains for 24 hours. *n*=3-9 biological replicates with 3-6 technical replicates each. Significance determined by one-way ANOVA with Dunnett’s multiple comparisons test. \*\*\*\**P*<0.0001. (B) Fluorescence microscopy images from mutant screens shown in A. Images are representative of 3 biological replicates with 3-6 technical replicates each. Scale = 80µm.

Hypothesizing that an agr-regulated secreted enzyme was responsible, we next tested CFCM from several transposon mutants with disruptions in genes encoding agr-regulated proteases, which are known to play a role in biofilm dispersal in *S. aureus* (24), and lipases, which we believed may alter the corynemycolic acids of the *Corynebacterium* cell wall (Fig 5A; red bars). However, these mutants all retained anti-aggregation activity similar to the wild-type *Sa* CFCM. We next tested mutants with transposon insertions in genes encoding phenol-soluble modulin (PSM) transporter proteins, *pmtB* and *pmtC*, as PSMs are known to be heat-resistant, agr-regulated proteins (11, 25–27). The *pmtB* transposon insertion partially restored *C. pseudodiphtheriticum* aggregation, and CFCM from the *pmtC* transposon insertion significantly abrogated the inhibitory effects of *S. aureus* CFCM (Fig 5A; gray bars). To further narrow down which PSMs were responsible, CFCM from a *S. aureus psm*α1-4 deletion strain, lacking ⍺-type PSMs 1 through 4, and a *psm*α1-4 deletion encoding an additional start codon mutation (ATG>ATT) in the *hld* gene, encoding the 5^th^ ⍺-type PSM, δ-hemolysin, was tested (28). Only the *psm*α1-4 *hld*^ATT^ double mutant lost the ability to inhibit *C. pseudodiphtheriticum* aggregation (Fig 5; gray bars).

To confirm results observed with CFCM from *S. aureus* mutants, we next examined whether addition of recombinant PSM peptides could inhibit *C. pseudodiphtheriticum* aggregation (Fig. 6). Production of the different types of PSMs encoded by *S. aureus* has been shown to vary substantially (29), therefore we tested addition of δ-toxin and PSMα3 peptides individually. Addition of purified, recombinant PSM peptides ranging from 0.6 to 5 µg/mL significantly inhibited *C. pseudodiphtheriticum* aggregate formation for both recombinant δ-toxin and PSMα3 in a dose-dependent manner (Fig. 6AB). Peptide concentrations evaluated were calculated to be equivalent to concentrations that would be present in up to 10% v/v *Sa* CFCM, based on previous measurements of PSMs generated in overnight stationary-phase *S. aureus* cultures (29). Neither recombinant δ-toxin nor PSMα3 affected *C. pseudodiphtheriticum* growth (Fig. 6C). We then tested whether δ-toxin is capable of dispersing pre-formed *C. pseudodiphtheriticum* aggregates using time-lapse microscopy (Fig. 6D; Movie S2). Compared to the untreated control, *Corynebacterium* aggregates were disassembled by the addition of recombinant ∂-toxin, with substantial disruption occurring within two hours.

**Figure 6:**
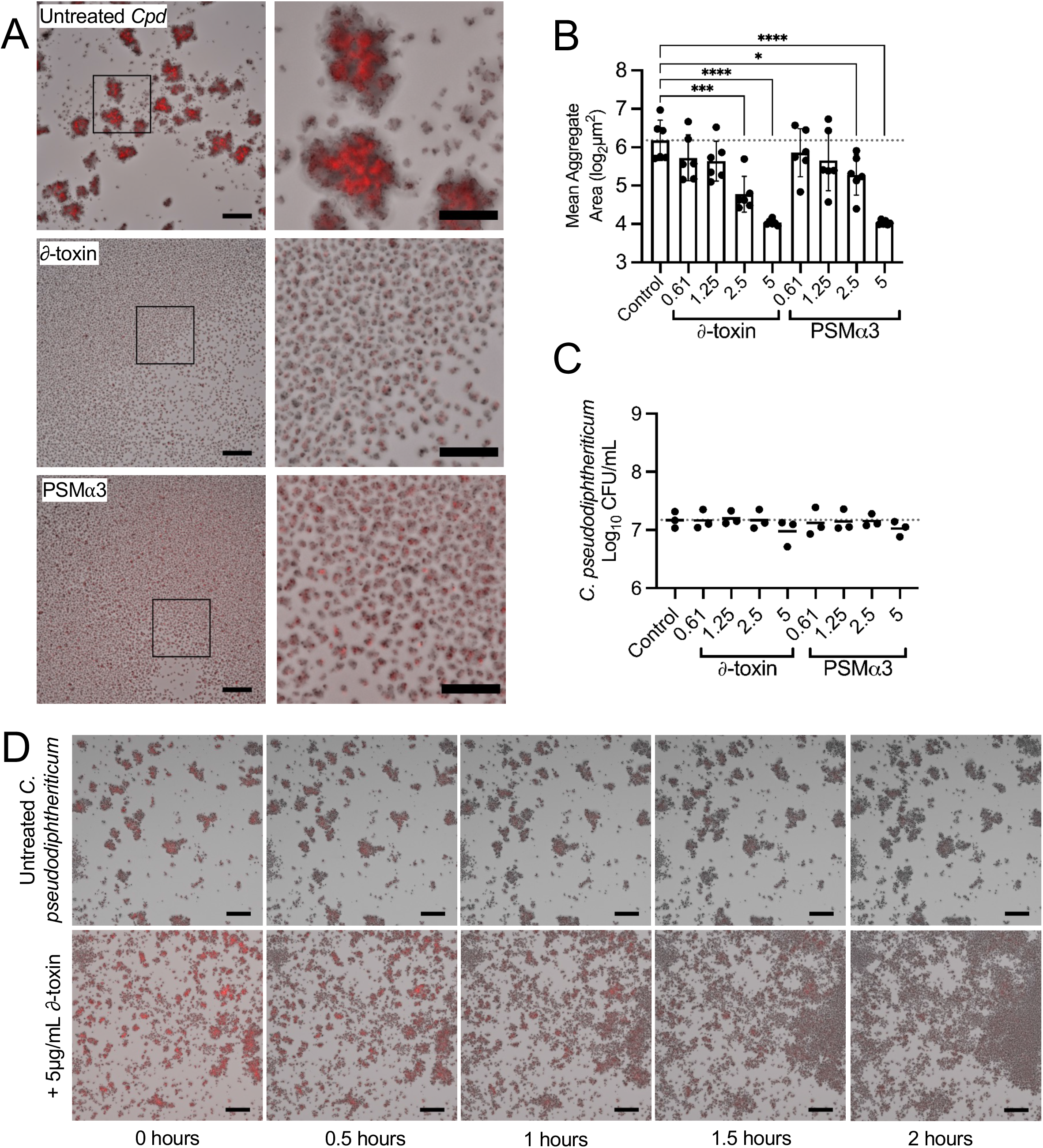
*S. aureus* phenol-soluble modulins inhibit *Corynebacterium pseudodiphtheriticum* aggregation and induce dispersal. (A) tdTomato *C. pseudodiphtheriticum* grown for 20 hours with 5µg/mL recombinant δ-toxin or PSMα3. Insert shown on right. Images are representative of 3 biological replicates with 3 technical replicates each. (B) Mean aggregate area quantification of microscopy images as shown in A. *n*=6 biological replicates with 3-6 technical replicates each. Significance determined by one-way ANOVA with Dunnett’s multiple comparisons test. \**P*<0.05 and \*\*\*\**P*<0.0001. (C) Colony-forming units of *C. pseudodiphtheriticum* for A,B. *n*=3 biological replicates with 3 technical replicates each. (D) Timelapse images of *C. pseudodiphtheriticum* exposed to 5µg/mL recombinant δ-toxin after 20 hours of growth. Images are representative of 3 biological replicates with 2 technical replicates each. Scale = 80µm, 40µm for 8X magnification

### *Staphylococcus aureus* phenol-soluble modulins alter *Corynebacterium pseudodiphtheriticum* adherence to nasal epithelial cells

We previously observed *Corynebacterium* CRS clinical isolates could colonize and form large aggregates in a co-culture model using polarized, air-liquid interface human nasal epithelial cell (HNEC) cultures (21). Given our observations of the effects of *Sa* secreted factors on *Corynebacterium* aggregation in vitro, we next asked if *Sa* CFCM would be sufficient to limit *C. pseudodiphtheriticum* growth in association with HNECs. We observed addition of CFCM from WT *S. aureus* strains resulted in a significant reduction in *C. pseudodiphtheriticum* adherence to HNECs at 1 hour with WT CFCM, but not CFCM from the *agrA::Tn* strain (Fig. 7A), supporting earlier our observations in the *in vitro* aggregation assay (Fig. 5). To examine whether *S. aureus* PSMs could interfere with *C. pseudodiphtheriticum* adherence and aggregate formation on HNECs, we inoculated *C. pseudodiphtheriticum* in the presence or absence of recombinant δ-toxin. Co-incubation of *C. pseudodiphtheriticum* with recombinant δ-toxin was sufficient to significantly decrease adherence at 1-hour (Fig. 7B). To confirm whether this effect from recombinant δ-toxin was targeting the *Corynebacterium* more-so than the HNECs, 1-hour adherence to HNECs was examined after a pre-incubation of either the *Corynebacterium* or the HNECs with 5µg/mL δ-toxin for one hour (Fig 7C). Comparing these groups showed a significant decrease in adherence with the pre-incubated *Corynebacterium* over the pre-incubated HNECs. Extending the co-incubation with δ-toxin to 6 hours resulted in a significant decrease in *C. pseudodiphtheriticum* colonization of HNECs (Fig. 7D-F). Fluorescence microscopy of tdTomato-labeled *C. pseudodiphtheriticum* colonizing HNECs in the same conditions showed marked reduction in formation of *C. pseudodiphtheriticum* aggregates in the presence of δ-toxin with lower total biomass as quantified through NIS-elements (Fig. 7D,E). Quantifying colony-forming units for *C. pseudodiphtheriticum* at the 6-hour timepoint demonstrated a significant decrease in *Corynebacterium* colonizing the HNECs when grown in the presence of δ-toxin (Fig 7F).

**Figure 7:**
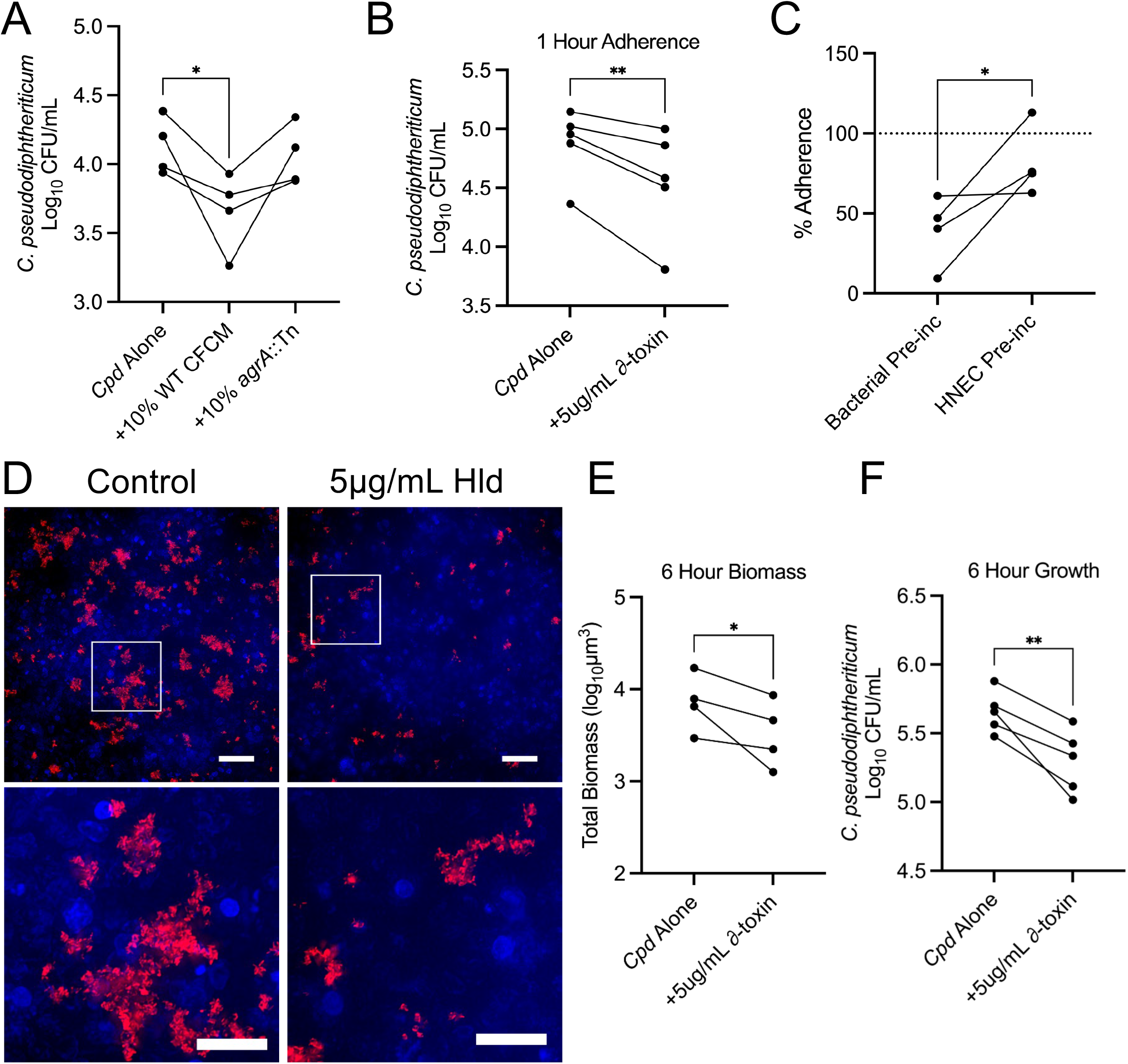
∂-toxin reduces *Corynebacterium pseudodiphtheriticum* adherence to nasal epithelial cells. (A) Colony-forming units of *C. pseudodiphtheriticum* attached to human nasal epithelial cells after a 1-hour co-incubation with 10% *S. aureus* CFCM. *n*=4 matched biological replicates. Significance determined by one-way ANOVA with Dunnett’s multiple comparisons test. \**P*<0.05. (B) Colony-forming units of *C. pseudodiphtheriticum* co-incubated with δ-toxin on HNECs for 1 hour. *n*=5 matched biological replicates. One-tailed paired t-test. *\*\*P*<0.01. (C) Adherence of *C. pseudodiphtheriticum* to HNECs following a 1-hour pre-incubation of 5µg/mL δ-toxin with the bacteria or HNECs. Data is normalized to the untreated control denoted by dashed line. One-tailed t-test \**P*<0.05. *n*=4 matched biological replicates. (D) Microscopy images (40X magnification; highlighted region shown below) of tdTomato-labeled *C. pseudodiphtheriticum* adherent to HNECs after a 6-hour co-incubation with 5µg/mL δ-toxin. Scale bar = 40µm; 20µm for 8X magnified region. (E) Biomass volume quantification of images from D. One-tailed paired t-test *\*P*<0.05. (F) Colony-forming units of *C. pseudodiphtheriticum* attached to HNECs after 1-hour with a 1-hour pretreatment of the bacteria or HNECs with 5µg/mL δ-toxin. Two-tailed t-test \**P*<0.05. (F) Colony-forming units of *C. pseudodiphtheriticum* on HNECs after 6 hours of growth with or without 5µg/mL δ-toxin. *n*=5 matched biological replicates. One-tailed t-test \*\**P*<0.01.

## DISCUSSION

The healthy upper respiratory tract harbors numerous bacterial species that compete against each other through a variety of different mechanisms (18). In a diseased state, URT microbial diversity is significantly reduced, and the resulting dominance established by pathogenic organisms is a significant contributor to disease pathogenesis (30). While the detrimental role of *S. aureus* in URT disease is well established, and previous work has identified many potential polymicrobial interactions that may occur between *S. aureus* and other URT colonizers (17, 18), few studies to date have assessed how *S. aureus* might mediate a transition from the healthy, microbially diverse URT to a diseased, pathogen-dominated URT via direct competition with URT commensal competitors who occupy the same niche. Here, we demonstrate a novel role for *S. aureus* ∂-toxin, a CRS-associated virulence factor (7), in altering the commensal lifestyle and commensal-host interactions for a competing URT commensal colonizer, *Corynebacterium pseudodiphtheriticum*.

In the data we present here, we demonstrate the agr-regulated *S. aureus* toxins ∂-toxin and PSMα3 can prevent *Corynebacterium* aggregation, disperse pre-formed aggregates, and limit colonization of host nasal cells. Previous studies of *Corynebacterium* and *S. aureus* interactions showed that *Corynebacterium* can alter *S. aureus* behavior through inhibition of agr quorum sensing or by inducing killing in *S. aureus* strains with an active agr system (15, 16). This partially explains the negative correlation observed between *Corynebacterium* and *S. aureus* in the URT (18–20, 31). However, when *S. aureus* does not have a functional agr system, its resulting gene expression is commensal-like and does not diminish its ability to colonize the upper respiratory tract, as numerous studies have demonstrated (11, 12, 15). Further, *S. aureus* populations in the URT can be heterogeneous, with a portion being incapable of agr activation and therefore immune to *Corynebacterium*-induced killing (32). The ability of *Corynebacterium* species to inhibit or exploit the *S. aureus* agr system suggests that agr activation likely exerts a selective pressure on *Corynebacterium* co-existing with *S. aureus* in the URT. Biofilms and biofilm-like lifestyles, such as aggregation exhibited by *Corynebacterium*, are vital for some colonizing species (33). The ability of *S. aureus* to disrupt this aggregation lifestyle in *Corynebacterium* (Fig 1) could be the evolutionary pressure needed to conserve a method by which to inhibit *S. aureus*, specifically through the agr system that governs production of PSM toxins. Further supporting this, we found that a functional agr system is necessary for antagonism of *Corynebacterium* aggregation (Fig 5). We saw that aggregation of three separate species of *Corynebacterium* (Fig 3) was strongly inhibited by *S. aureus* secreted factors, suggesting a broadly antagonistic relationship between commensal *Corynebacterium* species and *S. aureus*.

The significant association between *S. aureus* agr-regulated toxins and CRS pathogenesis is strong evidence that agr activation occurs in the URT in inflammatory disease and results in production of ∂-toxin (5, 10, 34, 35). Indeed, proteomics analysis of mucosal tissue samples from CRS patients found ∂-toxin and multiple agr-related proteins (7). Our observation that agr-regulated ∂-toxin can significantly disrupt *Corynebacterium* aggregation (Fig. 5,6) and its ability to colonize nasal epithelial cells (Fig 7) offer a possible mechanism by which *S. aureus* can create not only an inflammatory environment, but also an environment that is inhospitable for *Corynebacterium* and other commensal microbes reliant on aggregation or biofilm formation. As dysbiosis is a major component of CRS disease, (30) and multiple URT commensals have mechanisms for antagonizing *S. aureus* (17, 36), it is reasonable that *S. aureus* is actively contributing to this process in an effort to outcompete other microorganisms. Supporting this, a recent study examined gene presence for agr-regulated toxins in *S. aureus* and found that while presence of most genes ranged widely amongst CRS isolates, *hld* was conserved in 100% of isolates evaluated (37). Additionally, hypoxic conditions can increase agr activation and PSM production (38, 39), and the occluded sinuses in CRS are known to be a microoxic environment (6, 40, 41). Further, a prior study observed variable levels of *S. aureus* agr activation in the presence of representative sinus microbial communities cultured anaerobically, with cell-free supernatants from some communities leading to heightened agr activation (40). While *hld* is widely conserved, production of ∂-toxin can vary greatly between strains (42). Our own data supports this, as multiple *S. aureus* strains were not capable of preventing *Corynebacterium* aggregation, specifically Mu50 and N315 (Fig 2). This may be explained by the lack of ∂-toxin production by Mu50 and low production by N315 in a study by Su and colleagues (42). In this same study, production of ∂-toxin by *S. aureus* strain 502A was well-above the average, consistent with our finding that 502A CFCM prevents *Corynebacterium* aggregation (Fig 2). Although direct quantification of PSMs in the URT has not occurred been done, a recent study measured PSM production in a murine air pouch infection model (29). A 1 mL lavage of the air pouch after 48 hours of infection contained on average >1µg/mL ∂-toxin, which supports that *S. aureus* is capable of producing high concentrations of ∂-toxin during infection, consistent with the significantly higher quantities of ∂-toxin and low levels of other α-type PSMs in *S. aureus* CFCM (29).

The complex role of ∂-toxin in *S. aureus* biology limits the conclusions that can be drawn from CFCM studies. PSMs are known to regulate *S. aureus* biofilm structure, mediating detachment and microchannel formation, and PSM production was variable depending on the biofilm microenvironment (43). The dispersal ability of PSMs on *S. aureus* biofilms is believed to be through their proposed activity as surfactants that can mediate spreading on wet surfaces (44). ∂-toxin is believed to play a key role in the formation of *S. aureus* extracellular vesicles, which can contain numerous *S. aureus* products, as well as in amyloid fiber formation (29, 45). However, the other ⍺-type PSMs have been implicated in vesicle formation (46). Additionally, amyloid fibers have been found to be resistant to protease treatment (29), and the tertiary structure of some PSM amyloid fibers are sensitive to heat treatment (45). Our results shown in Figure 4 implicate sensitivity to protease treatment and resistance to heat, suggesting that amyloid fibers are not required for aggregation inhibition of *C. pseudodiphtheriticum*. Although our findings implicate *hld* being necessary for aggregation inhibition (Fig 5), addition of recombinant PSMα3 was sufficient to inhibit aggregate formation in *Corynebacterium* (Fig 6). We predict the PSMα knockout strain used in Figure 2 retained the ability to inhibit *C. pseudodiphtheriticum* aggregation due to the significantly lower production of PSMα3 compared to ∂-toxin that has been previously reported (29). Our data suggest a conserved ability of alpha-type PSMs to inhibit *Corynebacterium* aggregation, which may stem from its previously reported surfactant-like activity (44). Although a previous study found bactericidal effects for proteolytic derivatives of PSMs (47), we did not observe significant growth limitation by *S. aureus* secretions (Fig 1) and no such effect for either full length PSM peptide at the highest levels tested (Fig 6). Bactericidal effects from PSM production may be limited to certain species, with the possibility that outer membranes in Gram-negative and some Gram-positive bacteria offering a protective effect against PSM derivatives. Future studies will examine the mechanisms behind ∂-toxin inhibition of *Corynebacterium* aggregation and ask if this anti-aggregation effect is specific for competition against *Corynebacterium* species or broadly effective against other URT colonizing bacteria.

Collectively, these experiments lead us to hypothesize that *S. aureus* is capable of directly interfering with the *Corynebacterium* commensal lifestyle in the URT through production of ∂-toxin and alpha PSMs, contributing to the reduction in *Corynebacterium* abundance often observed in the diseased CRS microbiome. Several observations made by other groups support this hypothesis. *S. aureus* commensalism is linked to the inactivation of the agr system, either through genetic mutations, inhibition by *S. aureus* gene regulators, or external inhibition from competing bacteria (15–18). During or prior to *S. aureus*-associated CRS pathogenesis, agr is activated, leading to the secretion of several virulence factors, including ∂-toxin, that have been found in tissue samples obtained from subjects with CRS (7, 10, 35, 48–50). We speculate that inhibition of *Corynebacterium* aggregation by ∂-toxin provided the evolutionary pressure for *Corynebacterium* species to develop a yet unidentified method of inhibiting *S. aureus* agr activation. The ability of ∂-toxin to reduce adherence of *C. pseudodiphtheriticum* to nasal epithelial cells and disperse pre-formed aggregates may promote immune clearance of *Corynebacterium* species and other commensal colonizers, as well as increase susceptibility to antibiotics, which will be investigated in future studies.

## MATERIALS AND METHODS

### Bacterial strains and culture

*Staphylococcus aureus* and *Corynebacterium* spp. strains used in this work are detailed in Table S1. Bacterial stocks were stored at −80°C in 20% glycerol. Bacteria from frozen stocks were streaked onto brain-heart infusion (BHI) agar plates, with antibiotics if containing selective markers. Overnight cultures were inoculated with a single colony from agar plates. BHI broth was used for in vitro culture of all strains. *Corynebacterium* spp. cultures were incubated at 30°C with shaking at 200 RPM, while *S. aureus* strains were incubated at 37°C with shaking for 24 hours. *C. accolens* culture media was supplemented with 0.05% Tween 80 for overnight cultures. Kanamycin (50µg/mL, ThermoFisher) was added to tdTomato-expressing *C. pseudodiphtheriticum* agar plates and cultures for maintenance of the pJOE7706.1-tdTomato plasmid. Transposon insertion mutants from the Nebraska Transposon Mutant Library were grown with 50µg/mL erythromycin (ThermoFisher) on BHI agar. Transposon mutant strains from the Network on Antimicrobial Resistance in *Staphylococcus aureus* (NARSA) and *C. amycolatum* were obtained from BEI Resources, NIAID, NIH.

### Aggregation inhibition assay

*S. aureus* strains inoculated into 5mL BHI broth were incubated for 24 hours at 37°C with shaking at 200 RPM. Cultures were centrifuged at 3,000 RCF for 5 minutes to pellet bacteria. Culture supernatant was filter sterilized using a 0.22µm-pore syringe filter (Fisher Scientific) to obtain *S. aureus* cell-free conditioned media (CFCM). *C. pseudodiphtheriticum* with pJOE7706.1-tdTomato plasmid was diluted into BHI broth to an A_600_ of 0.002 along with 0.2mM isopropyl β-d-1-thiogalactopyranoside (IPTG; Sigma Aldrich) to induce tdTomato expression and 100µg/mL kanamycin. CFCM from *S. aureus* was diluted with BHI broth to 2X concentration of the desired v/v percentage. 50µL of the cell-free conditioned medium was added to an untreated 96-well microtiter dish (Corning) followed by 50µL of 2X *C. pseudodiphtheriticum*/IPTG mixture and kanamycin. The plate was incubated at 30°C without shaking for 20 hours. For other strains of *Corynebacterium*, the assay was performed without kanamycin or IPTG, with 16 hours of growth in BHI broth. Imaging was performed using a Nikon Eclipse Ti2 widefield using a Plan Apo VC 20X Air N2 lens. Images were acquired from the center of each well (±300µm) to limit variation of brightfield intensity due to edge effect.

### Image analysis and quantification

Analysis of images used the Nikon NIS-Elements AR software package (version 5.42.02 Build 1801). Quantification of aggregate size was determined using the Object Counting tool in NIS-Elements. Minimum intensity projections of the z-stack were created followed by auto-thresholding of the brightfield intensity using the Otsu Original method. Raw data from the object counting tool was exported as a tab delimited text file prior to analysis. Text files were imported into RStudio (version 2024.04.0 Build 735 “Chocolate Cosmos” Release). Area values below 1 we excluded to reduce exceedingly small values and to allow for log_2_ transformation. Mean aggregate area for an individual well was calculated and averaged amongst wells for that group using base R.

### Live imaging microscopy

Timelapse microscopy was performed under similar conditions as listed above with the following modifications. Glass bottom Ibidi µ-Slide dishes were used for compatibility with the Nikon Perfect Focus System. Due to the increased surface area of the wells, 300µL was the final volume as opposed to 100µL. For dispersal with recombinant peptides, *C. pseudodiptheriticum* was grown for 20 hours prior to addition of synthetized peptide or phosphate-buffered saline serving as the vehicle control. Incubation continued for 4 hours after addition of the synthetic peptide.

### Cell-Free Conditioned Medium (CFCM) Treatments

Heat treatment as well as proteinase-K incubation/inactivation utilized thermocyclers for temperature consistency and replicability. BHI broth, serving as a treatment control and *S. aureus* cell-free conditioned medium (100% v/v), was heated to 98°C for 50 minutes followed by 10 minutes at 4°C. Protease treatment used 100µg/mL proteinase-K (New England Biosciences) with a 50-minute incubation at 56°C followed by a 10-minute heat-inactivation step at 98°C. For size fractionation, Amicon 3kDa cutoff centrifugal filters (Millipore-Sigma) were used. Flow through was used directly in the assay whereas the concentrate was diluted in fresh BHI to the original volume followed by centrifugation at 13,000 RCF for removal on precipitates.

### Human nasal epithelial cell co-culture assays

Cell culture experiments were conducted as described previously with slight modifications (21). The RPMI 2650 human nasal epithelial cell line obtained from ATCC (ATCC CCL-30) was cultured in Eagle’s minimal essential medium (MEM; Corning) supplemented with L-glutamine (Gibco), 10% fetal bovine serum (Gibco), Pen-Strep (Gibco) and Plasmocin (Invivogen). Trypsin (Corning) was used to lift nasal cells prior to seeding in 6.5mm transwell filters (CellTreat) pre-coated with Vitrogen plating medium as described previously (21). Transwell filters were incubated for one week prior to removing apical media for the transition to air-liquid interface. Nasal cells were then maintained at air-liquid interface for one week prior to use in co-culture assays. Colonization with *C. pseudodiphtheriticum* was tested by inoculating the apical surface of ALI cultures with bacteria at an A_600_ of 0.1 in serum- and antibiotic-free MEM. For colony-forming unit assays, a 100µL wash with MEM was used to removed non-adherent bacteria from the cell surface followed by 15 minutes of orbital shaking at 200rpm with 100µL of MEM with 0.1% Triton X-100. Nasal cells were then scraped from the surface of the transwell filters and briefly vortexed to ensure dissociation of bacteria prior to serial dilution and viable colony forming unit counts. For microscopy, transwell filters were fixed with 4% paraformaldehyde (Electron Microscopy Sciences) overnight at 4°C. Filters were then washed with Dulbecco’s phosphate-buffered saline (DPBS; Sigma Aldrich) followed by permeabilization in PBS with 0.1% Triton X-100 (Biorad). DNA was stained with Hoechst 33342 (Invitrogen) prior to mounting with ProLong Gold (Invitrogen). Fluorescence microscopy was performed using a Plan Fluor 40X Oil DIC H N2 lens with Type B non-drying immersion oil (Cargille Laboratories on a Nikon Eclipse Ti2 microscope as described above.

For assays evaluating *S. aureus* CFCM, 10% v/v CFCM was added concurrently with *C. pseudodiphtheriticum* inoculums for adherence assays. Viable colony forming units (CFUs) were collected and enumerated on BHI agar plates after 1 hour or 6 hours of co-culture with HNECs as indicated. For assays evaluating the effects of recombinant PSM peptides, *C. pseudodiphtheriticum* was inoculated with 5ug/mL ∂-toxin or vehicle control.

### Recombinant peptide assays

Recombinant PSM⍺3 and ∂-toxin (IBT Bioservices) were stored at −80°C. Prior to use in assays, the peptides were thawed in a 37°C water bath. Peptides were gently mixed and diluted into BHI broth for aggregation assays or MEM for HNEC co-culture assays at a 2X concentration. Control groups for these assays received an equal volume of vehicle, either MilliQ water for PSM⍺3 or phosphate-buffered saline for ∂-toxin.

### Statistical analysis

GraphPad Prism (GraphPad Software; Version 10.2.3 Build 347) was used for statistical analyses of colony-forming unit assays and aggregation assays. Normality of data was assessed with QQ-plots, probability of Gaussian distribution, and Shapiro-Wilk’s test. For one-way ANOVA tests, Brown-Forsythe test was used to evaluate homogeneity of variance and Shapiro-Wilk’s test to check normality of residuals. Dunnett’s multiple comparisons test was used for post-hoc analysis. For paired t-tests, Shapiro-Wilk’s test for verifying normality of residuals. Error bars denote standard deviation. Lines shown on graphs for individual groups represent the mean.

## Acknowledgments

We thank Dominique Limoli and Alex Horswill for sharing PSM knockout strains. This work was supported by 5T32GM008111-35 to J.T.H. and CFFROWE21R3 to M.R.K. The funders had no role in study design, data collection and interpretation, or the decision to submit the work for publication.

## Author Contributions

JH and MK performed or contributed to experiments that generated data for this manuscript. JH and MK analyzed data and wrote the manuscript.

