## Supporting Information text file for "*Staphylococcus aureus* Phenol-Soluble Modulins Mediate Interspecies Competition with Upper Respiratory Commensal Bacteria"

**SUPPLEMENTAL FIGURE LEGENDS**

**Figure S1: *Corynebacterium pseudodiphtheriticum* aggregate size increases over time.** (A) Timelapse of tdTomato C*. pseudodiphtheriticum* grown over 20 hours in brain-heart infusion broth. 8X magnification of inset on bottom row. Scale bar = 80µm; 40µm for 8X magnification. (B) Quantification of mean aggregate size for A. Lines indicate a linear regression for each biological replicate. n=3 biological replicates with 6 technical replicates each. Slopes were not statistically different (P=0.2072).

**Movie S1: *Corynebacterium pseudodiphtheriticum* grows in aggregate form. (**A) tdTomato *C. pseudodiphtheriticum* grown for 20 hours in brain-heart infusion broth.

**Movie S2:** ***S. aureus* phenol-soluble modulins directly inhibit and disperse *Corynebacterium pseudodiphtheriticum* aggregation.** Timelapse of *C. pseudodiphtheriticum* culture exposed to δ-toxin after 20 hours of growth.

SUPPLEMENTAL TABLES

Table S1. Bacterial Strains

| **Strain name** | **Description** | **Reference** |
| --- | --- | --- |
| ***Staphylococcus aureus* strains** | | |
| *S. aureus* LAC 13c | USA300 CA-MRSA, erm^S^ | (1) |
| *S. aureus* USA100 | MRSA CC5 isolate | (2) |
| *S. aureus* SH1000 | sigB+ derivative of NCTC8325-4 | (3) |
| *S. aureus* 502A | Human nasal colonizing isolate | (4) |
| *S. aureus* Newman | MSSA, Type 5 capsule producer | (5) |
| *S. aureus* N315 | Hospital-acquired MRSA | (6) |
| *S. aureus* Mu50 | Hospital-acquired MRSA | (6) |
| *S. aureus* SF8300 | USA300 MRSA wound isolate | (7) |
| *S. aureus* LAC 13c *psmα*- | USA300 LAC ∆psm⍺1-4 | (8) |
| *S. aureus* LAC 13c *psmα*-/*hld*^ATT^ | USA300 LAC Δ*psm*ɑ1-4 δATG-ATT | (8) |
| ***Corynebacterium*** **strains** | | |
| *Corynebacterium pseudodiphtheriticum* + pJOE7706.1-tdtomato | Sinus clinical isolate expressing pJOE7706.1 with tdtomato inserted at BsrGI and BamHI sites | (9) |
| *Corynebacterium propinquum* | Sinus clinical isolate |  |
| *Corynebacterium pseudodiphtheriticum* ATCC | ATCC strain 153, Lehmann and Neumann | (10) |
| *Corynebacterium amycolatum* | Strain SK46, skin isolate | (11) |
| **Nebraska Transposon Mutant Library *S. aureus* strains** | | |
| *S. aureus* *agrA::Tn* | Transposon Mutant NE1532 (SAUSA300_1992) | (1) |
| *S. aureus sigB::Tn* | Transposon Mutant NE1109 (SAUSA300_2022) | (1) |
| *S. aureus sarA::Tn* | Transposon Mutant NE1193 (SAUSA300_0605) | (1) |
| *S. aureus saeR::Tn* | Transposon Mutant NE1622 (SAUSA300_0691) | (1) |
| *S. aureus aur::Tn* | Transposon Mutant NE163 (SAUSA300_2572) | (1) |
| *S. aureus sspA::Tn* | Transposon Mutant NE1506 (SAUSA300_0951) | (1) |
| *S. aureus sspB::Tn* | Transposon Mutant NE934 (SAUSA300_0950) | (1) |
| *S. aureus splC::Tn* | Transposon Mutant NE1098 (SAUSA300_1756) | (1) |
| *S. aureus splF::Tn* | Transposon Mutant NE1764 (SAUSA300_1753) | (1) |
| *S. aureus geh::Tn* | Transposon Mutant NE1775 (SAUSA300_0320) | (1) |
| *S. aureus lip::Tn* | Transposon Mutant NE338 (SAUSA300_2603) | (1) |
| *S. aureus pmtB::Tn* | Transposon Mutant NE1188 (SAUSA300_1912) | (1) |
| *S. aureus pmtC::Tn* | Transposon Mutant NE1908 (SAUSA300_1911) | (1) |
