## Supplementary figures and images for "*Staphylococcus aureus* Phenol-Soluble Modulins Mediate Interspecies Competition with Upper Respiratory Commensal Bacteria"

### Figure S1

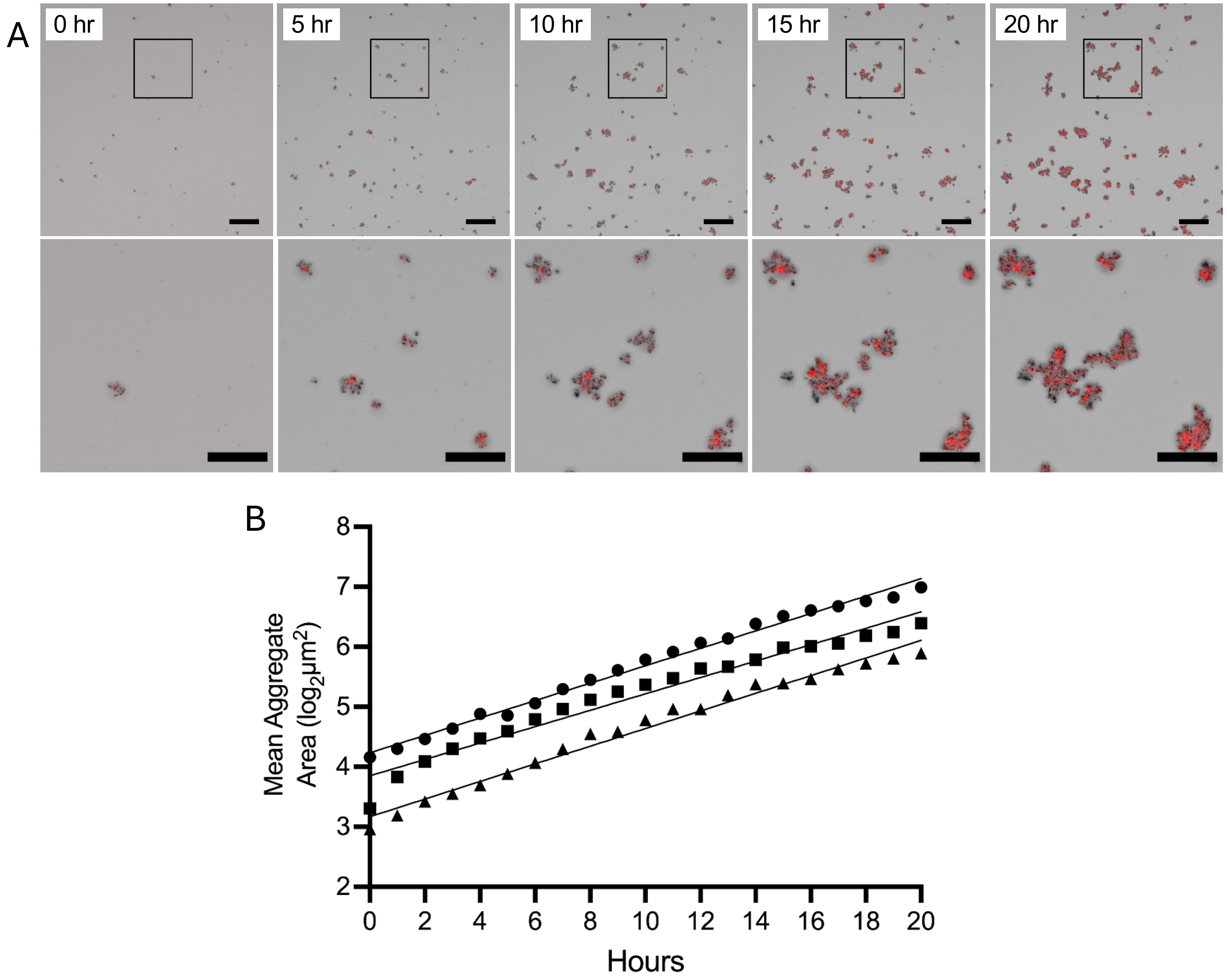
